# GN1, Glucosamine Analog, attenuate the neuronal hyperactivity of c-*fos* gene in Adjuvant-Induced Arthritis in Rats

**DOI:** 10.1101/2021.01.12.426455

**Authors:** Huma Jawed, Syed Uzair Ali Shah, Shahid I. Awan, Shazia Anjum, Shabana U. Simjee

## Abstract

The anti-inflammatory and anti-arthritic activity of GN1 (6-hydroxymethyl-3-(1-methylene-allylamino)-tetrahydro-pyran-2,4,5-triol), an analog of glucoseamine was investigated in the adjuvant induced arthritis (AIA) in rats. It was observed that the highly increased hind paw swelling was significantly reduced (p <0.001) with no noticeable retardation of body weight in the GN1 treated arthritic rats as compared to arthritic control rats. The histopathological analysis of isolated rat joints by scaling system classification of different stages appear in arthritic inflammatory lesion development, disclosed the suppressive effect in the inflammation processing in the GN1 treated animals. Different predictive markers such as TNF-α, nitric oxide IL-1β, peroxide and glutathione were also determined in the serum samples of the treated and non-treated arthritic rats in order to measure the intensity of inflammation. Our results show that GN-1 treatment in arthritic groups demonstrated marked reduction in *c-fos* expression. Results also revealed that GN1 significantly reduces the peroxide production (p < 0.002) and both the pro-inflammatory cytokines TNF-α (p < 0.015) and IL-1β (p < 0.001) in the arthritic rats receiving this treatment. Although, nitric oxide formation was found to be less than the arthritic control, but this reduction was not significant. The level of glutathione was significantly increased in GN1 treated arthritic rats (p < 0.037). These observations suggest that GN1 may help in reducing the inflammation and have anti-arthritic properties.

## INTRODUCTION

Chronic pain is a distressing and commonly prevalent problem among the world population. As compared to other medical conditions, patients suffered with chronic pain, experiencing effecting in their physical, psychological and social wellbeing (1–3). Arthritic pain is the second reported cause of pain in population (4). The disease is characterized by two phases, microvascular injury with synovial swelling and the influx of plasma components (5). The first phase is then followed by the accumulation of lymphocytes, macrophages and neutrophils in the joints. In chronic stages the cytology of joint show elevated leukocytes counts and numerous immune complexes. The subsequent interaction of these complexes with neutrophils initiates phagocytic responses and the release of inflammatory mediators such as TNF-α and IL-1β, proteolytic enzymes and reactive oxygen species (6). In addition, the proto-oncogene c-*fos* has also been proven to be induced in response to various inflammatory and nociceptive stimuli (24). The c-*fos* gene encodes protein c-Fos, a DNA-binding protein, which make transcriptional factor, activator protein-1 (AP-1), with c-Jun [19, 20]. In turn, C-Fos/C-Jun activate the gene expression of various pro-inflammatory signals including TNF-α and other cytokines. The activation of c-Fos expression in neurons can be used as a significant indicator of neuronal hyperactivity of central or peripheral inflammation and nociception [21–23, 24)

Various classes of drugs available including analgesic and anti-inflammatory for the suppression of rheumatoid arthritis (RA) by alleviating the arthritis associated symptoms and improve the quality of patient’s life (4). Despite the symptom’s suppression, none of available treatment are adequate therapy and cannot successfully reverse the arthritis pathogenesis. Among the available treatment, despite of the severe toxicities including, gastric ulcer and renal toxicities glucocorticoids and nonsteroidal anti-inflammatory drug (NSAIDs) are commonly prescribed for treatment of rheumatoid arthritis (7–13). Since then many compounds were introduced in search for drugs with anti-inflammatory activity. Now they are used in the combination with chondroprotective agents with disease modifying effects such as glycosaminoglycan, hyaluronan, glucoseamine, etc (14, 15). The use of glucoseamine has been widely reported to treat osteoarthritis (16, 17) and it was found to maintain the elasticity and flexibility of cartilage and connective tissues due to their ability to stimulate the biosynthesis of the glycosaminoglycans. Although glucoseamine is reported as anti-inflammatory agent (18, 19), but due to debilitating pain of joints it is not enough to relief it, and yet there is need to improve its biological activity. Thus, investigations for the new anti-inflammatory agents with minimum side effects are still a challenge and studies on both synthetic drugs and plants are conducted to realize this purpose.

In the present study, we have shown that GN1 (6-hydroxymethyl-3-(1-methylene-allylamino)-tetrahydro-pyran-2,4,5-triol), an analog of glucoseamine suppresses not only the major macroscopic changes such as swelling of hind and forelimbs but also inhibits the arthritic knee histopathologic changes. The efficacy of this compound was greater than indomethacin (5 mg/kg/day). In addition, the GN1 effectively attenuate the gene expression of c-*fos* and significantly decreased the levels proinflammatory cytokines (TNF-α and IL-1β) and reactive oxygen species (peroxide and nitric oxide) in the arthritic rats treated with this compound. Thus, GN1 is a potential active therapeutic agent for the treatment of rheumatoid arthritis and osteoarthritis. Our observations suggest that GN1 may help in reducing the inflammation and have anti-arthritic properties.

## METHODOLOGY

### Animals

For the experiments, Sprague Dawley (SD) female rats weighing 230-250 gm (8-10 weeks old) were used in this study. The experimental rats were kept at maintained temperature 21 ± 2°C and on light/dark cycle (12 hour each), standard laboratory rat food pellets and water were freely accessible to the animals. All the animals were randomly distributed to each treatment group.

### Induction of Arthritis

freshly prepared suspension (1mg/ 0.1mL) of lyophilize Mycobacterium tuberculosis MT37Ra (Difco Laboratories, USA) was intradermally injected once to induce arthritis in rats. Treatment of GN1 and Indomethacin were also started on the same day after induction of arthritis. (Stuart J. et al 1979).

### Clinical Assessment of Adjuvant Induced Arthritis (AIA)

On each alternate day, all the experimental rats in groups were evaluated for arthritis development by using following macroscopic scoring system (21, 22). The arthritis severity score for each rat was calculated by adding the scores of each individual paw.

| **Score** | **Observational score** |
| --- | --- |
| **0** | no signs of arthritis |
| **1** | redness / or swelling of the paw or 1 digit only |
| **2** | If 2 joints involved |
| **3** | If more than 2 joints involved |
| **4** | severe arthritis of the all digits and even entire paw |

### Measurement of Hind Edema

Clinical severity of arthritis was also determined by quantitating the change in the paw volume (as an indicator of edema) with a plethysmometer (model 7140, Ugo Basile, Varese, Italy). The advantage of using this method over diameter measurements of tibiotarsal joint is that it measures the limb in three-dimensions and therefore takes into account any variability of the pattern of swelling of individual limbs. The volume of a hind paw is reported as the mean ± SEM in ml. All measurements were made at the same time of day.

### Histological analysis

Rats were sacrificed at day 22; hind limbs were removed and fixed in 10% buffered formalin. The limbs were decalcified in 5% formic acid, processed for paraffin embedding, sectioned at 5-μm thickness, and subsequently stained with haematoxylin-eosin for examination under a light microscope (23). Sections were examined for the presence of hyperplasia of the synovium, pannus formation, and destruction of the joint space.

## REDOX ASSAY

The concentration of glutathione, peroxide and nitric oxide in serum were measured by a calorimetric assay kit, according to the manufacturer’s instructions (Quantichrome ™ Assay kit, Hayward, C.A., U.S.A.).

## MEASUREMENT OF CYTOKINES

The quantitative measurement of cytokine was done using ELISA assay kit as detailed in the guideline provided by the manufacturer (KOMA BIOTECH INC., Korea).

**Reverse transcriptase polymerase chain reaction (RT-PCR) analysis of c-fos expression**

### Primer for c-fos

The GAPDH (glyceraldehyde-3-phosphate dehydrogenase) was used as an internal standard (housekeeping gene). The primer sequence of the GAPDH gene and c-fos used in our study is shown in Table 2.

### Tissue collection and RNA extraction

At the end of the experiment, animals were anesthetized and transcardially perfused with 0.9 % saline. Brain tissue were removed and stored at −80 °C for subsequent RNA extraction. Brain regions including amygdala, hippocampus, thalamus, and cortex were isolated and subjected to homogenization. Proteinase K was added to the homogenate and kept at 55 °C for 10 min followed by centrifugation at 10,000 rpm for 3 min. The resultant supernatants were transferred into RNeasy spin column placed in a collection tube and re-centrifuged for 15 s at 10,000 rpm. The spin column membrane was washed twice for 15 s with the buffer at 10,000 rpm. To ensure that no ethanol is carried over during RNA elution, 500 μl buffer RPE was added in the spin column and centrifuged for further 2 min at 10,000 rpm. After washing, the RNeasy spin column is placed in a new 1.5-ml collection tube and 50 μl RNase free water was added directly to the spin column and centrifuged for 1 min at 10,000 rpm to elute the RNA. The quantification of the RNA was carried out by spectrometric analysis (Abs at 260 nm). An absorbance of 1 U at 260 nm corresponds to 40 lg of RNA per ml (A260 = 1) 40 lg/ml). The quantification of the total RNA was calculated as:

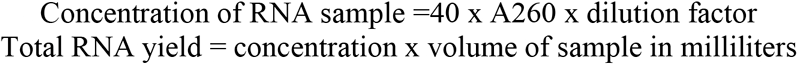

### cDNA synthesis and RT-PCR

The isolated RNA was reverse transcribed using Superscripts III First-Strand synthesis system (Invitrogen Corporation Carlsbad, CA, USA) for RT-PCR. Briefly, the components provided in the kit were combined in amounts described in the manufacturer’s protocol to make the RNA/ primer mixture and cDNA synthesis mix. The RNA/primer mixture was incubated at 65 °C for 5 min and then cDNA synthesis mix was added to this mixture and re-incubated at 50 °C for 50 min after which the reaction was terminated at 85 °C at a cycle of 5 min followed by placing the reaction contents on ice for 5 min. The final product i.e., cDNA was collected and stored at −20 °C. Negative control was also run and included RT reaction mixture omitting reverse transcriptase. The transcribed cDNA was amplified using Omniscript RT kit (Qiagen Inc.Valencia) and oligonucleotide primers corresponding to transcripts of the GAPDH and c-fos. The PCR mixture for each control reaction was prepared as outlined in the protocol. To exclude contamination of genomic DNA, reverse transcription was also carried out for the same sample without adding the enzyme (negative control). The PCR annealing temperatures employed are listed in Table 2. The products of reverse transcription reactions were placed in a preheated (94 °C) thermal cycler and then denatured for 1 min at 94 °C, followed by 35 cycles of amplification in the following manner: 15 s denaturation at 94 °C then 30 s annealing at 60 °C and then 1 min extension at 72 °C. The final extension step was performed at 72 °C for 10 min and upon completion the reactions were maintained on 4 °C. The PCR reaction product was electrophoretically resolved on 1 % agarose gel containing ethidium bromide. The loading dye (bromophenol blue) and ladder in 1:6 ratios was used to prepare working ladder. Samples and the ladder were then added to the valves and electrophoresed until the bromophenol is near the base of the gel. The bands on the gel were visualized under a UV lamp in a Gel-Dock System (Fluor Chem, Alpha INNOTECH).

### Statistical Analysis

The Statistical Package for the Social Sciences (SPSS) software was used to analyze the data. Throughout this study mean ± SE of means were used to describe the data in figures. The data were analyzed using two-way analysis of variance (ANOVA). The Bonferroni’s post hoc test was used to determine which group mean differs. With this test SPSS automatically adjusts the significant level for the multiple comparisons to avoid spurious significant differences being identified (any values below the level of 0.05 was considered as significant).

## RESULT AND DISCUSSION

In order to treat the symptoms and the pathological cause of the RA we have choices for glucocorticoids, non-steroidal anti-inflammatory agents and some biological agents having disease modifying abilities. Glucoseamine is one of the biological molecules that can be used in the RA to reconstruct the proteoglycan, restoring articular function and modify the symptoms of the disease. Beside its chondroprotective action and anti-inflammatory activity, there is need to improve these abilities because of the debilitating pain of the joint and bones in arthritis and glucoseamine alone is sometime not enough to overcome this. Therefore, our research team has synthesized the analog of glucoseamine i.e., GN1. To determine the macroscopically and biochemical effects of GN1 in the AIA rat model, the present study was designed. The development and progression of disease is assessed by mean arthritis severity score and paw volume (fig. 1-4). The signs arthritic score 3 in all groups were evident at day 10. In the non-treated arthritic group, the incidence was 100% at day 13 which remain persist at the end of the experiment. In contrast, the treatment with indomethacin, and GN1 exerted a significant attenuation in the incidence of AIA in GN1 treatment (P < 0.017) and indomethacin treatments (P < 0.01). Body weights was measured among the groups, in the initial days of the experiment there was normal increase in body weight of animal in all groups (fig. 3), but after day 10^th^ a significant decrease (P < 0.03) in body weight was observed in the arthritic rats receiving no treatment this loss continues until the end of the experiment. In contrast, the non-arthritic rats and arthritic rats treated with GN1 showed no further reduction in their body weight. From histopathological observation, GN1 was found to be suppressor of chronic cell infiltration, synovial hyperplasia and cartilage destruction (fig. 5).

**FIGURE 1:**
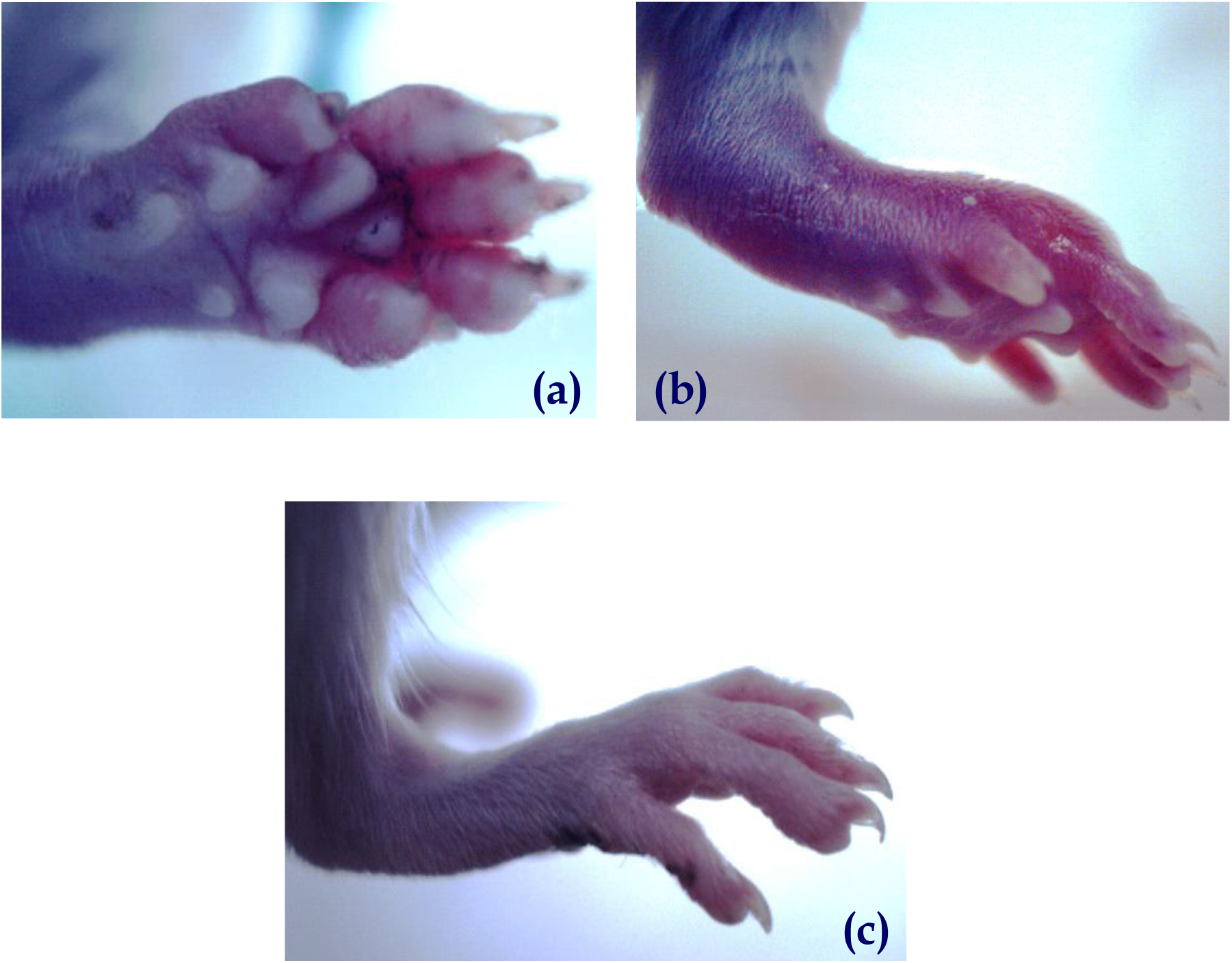
Onset of arthritic signs in the arthritic control group. The gradual increase in the erythema and inflammation is visible with the progression of the disease (a & b) as compared with the control group (c).

**FIGURE 2:**
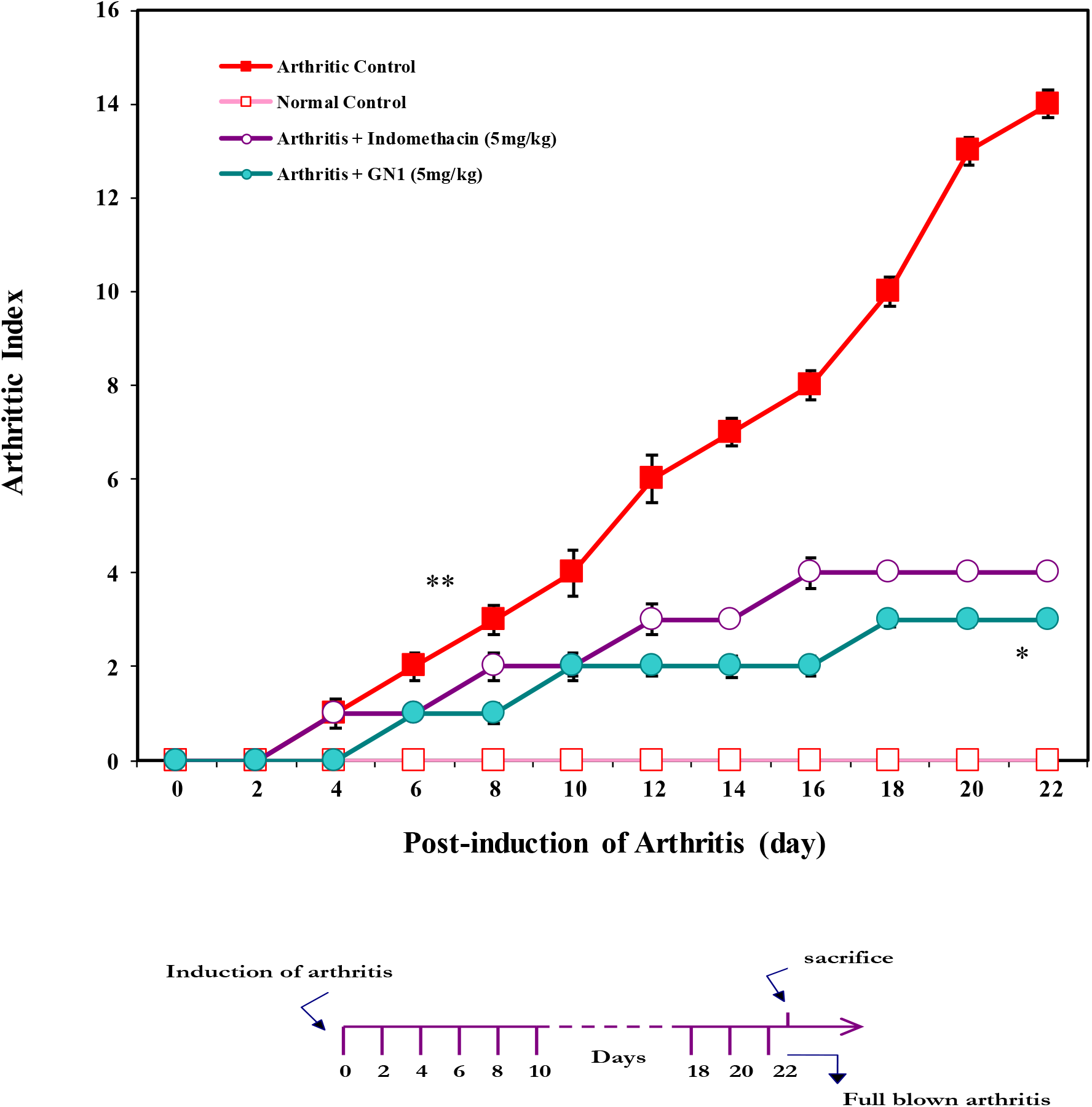
Arthritis Index (AI) of GN-1 treated arthritic animals. Each value represents the mean ± SEM of n = 12 animals per group. The data shows that the GN-1 treatment significantly reduces the arthritic index (*p < 0.017) compared to the arthritic control group. The arthritic control group exhibited a significant difference in their arthritic index from day 6 (**p < 0.001) onward as compared to the normal control group.

**FIGURE 3:**
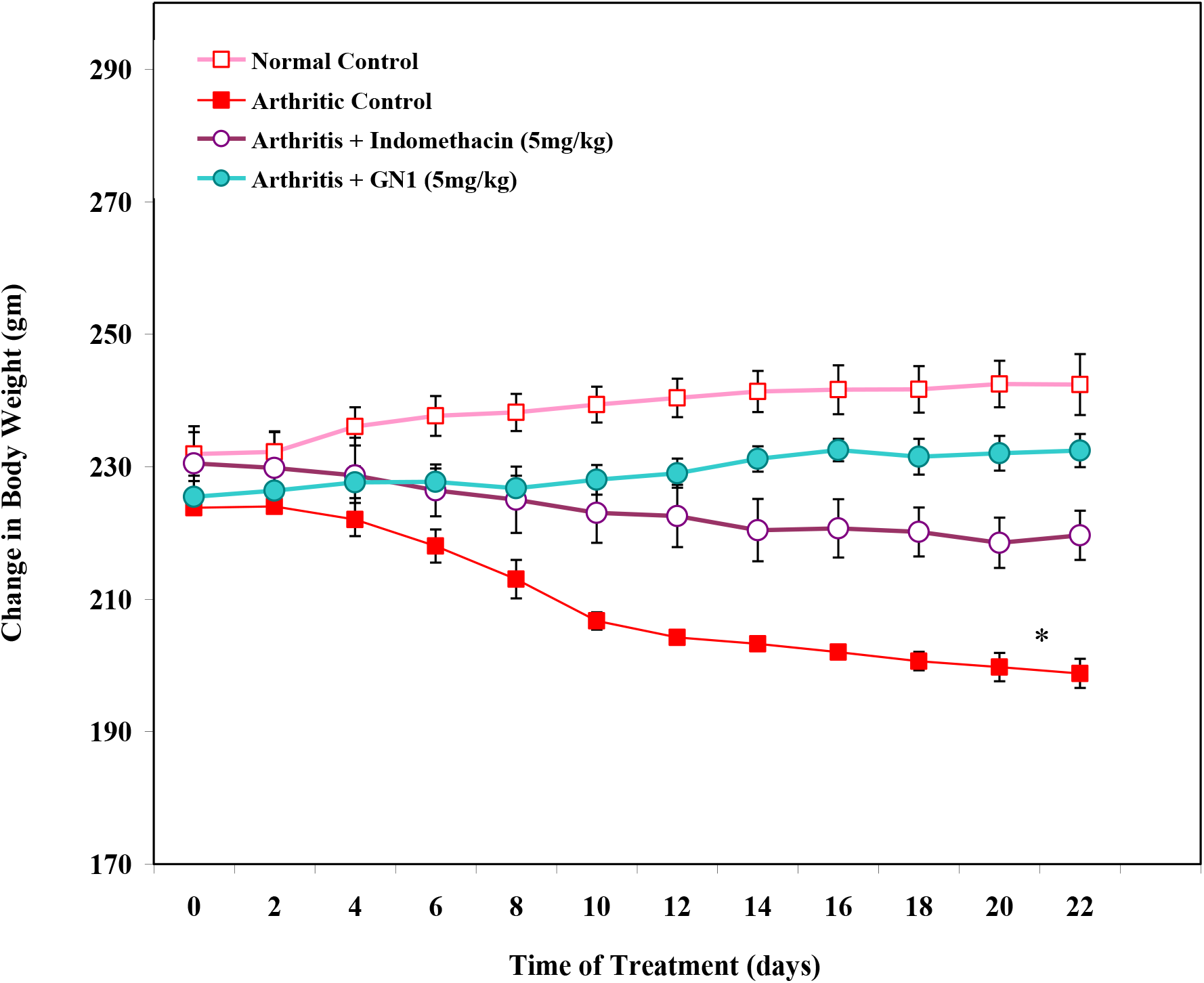
Change in body weight of GN1 treated arthritic rats over a period of 22 days post-induction of arthritis. A significant reduction (*p < 0.022) in the body weights of arthritic control animals was observed in comparison to normal control group. The loss of body weight was further observed in the arthritic control rats until the end of the experiment. In contrast, the arthritic rats treated with GN1 showed a slight but non-significant reduction in their body weights on day 8 but no further reduction was recorded in this group until the end of the experiment.

**FIGURE 4:**
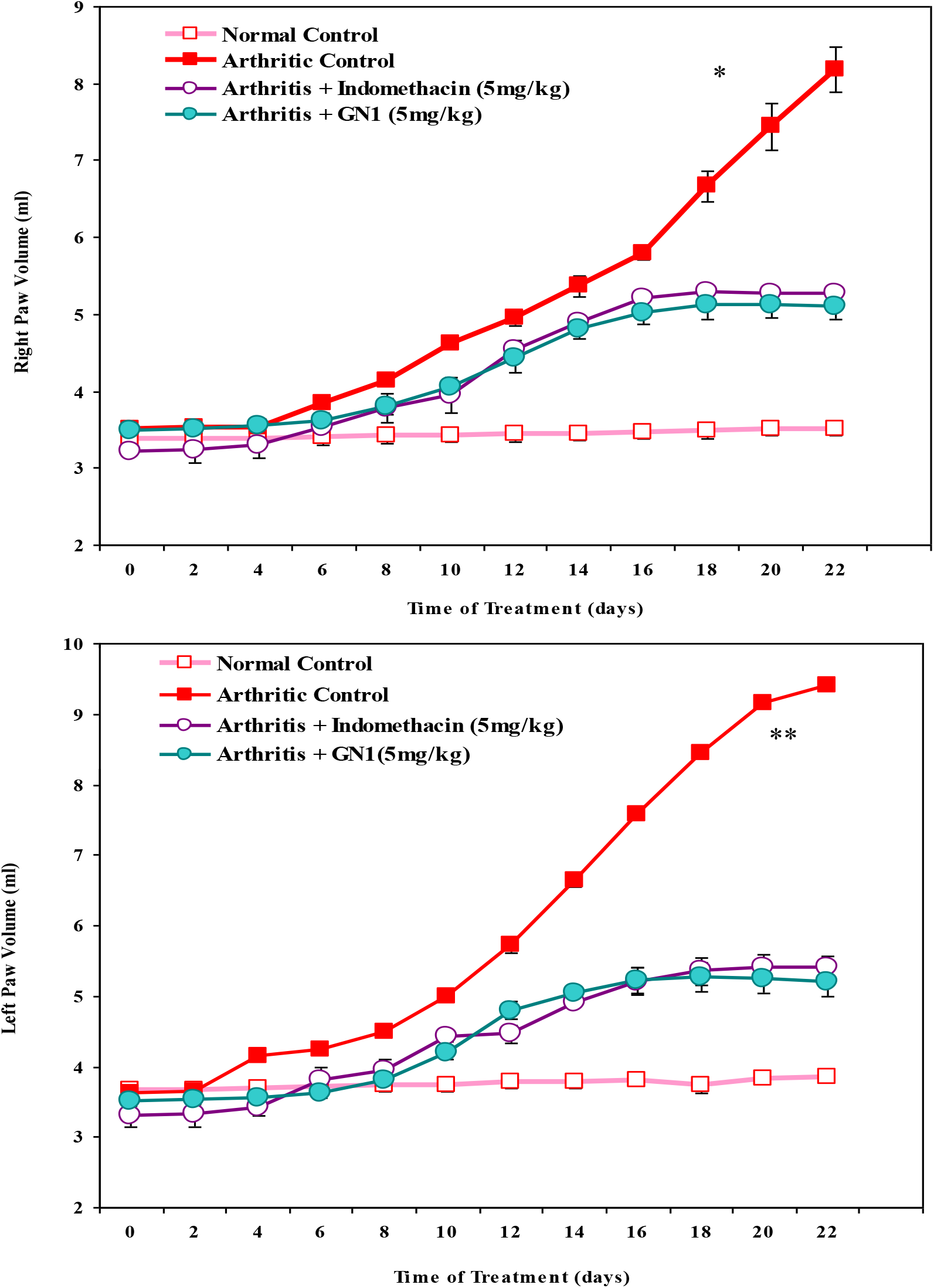
Effect of GN1 treatment on paw volume of arthritic rats. The increase in the paw volume of arthritic control group was found to be significantly different from the normal group on day 8, i.e., *p < 0.019 and **p < 0.004 for right and left paw respectively. In contrast, the arthritic animals treated with GN1 showed a gradual increase in their paw volumes from day 6 however, this increase was not progressed further from day 16 onward.

**FIGURE 5:**
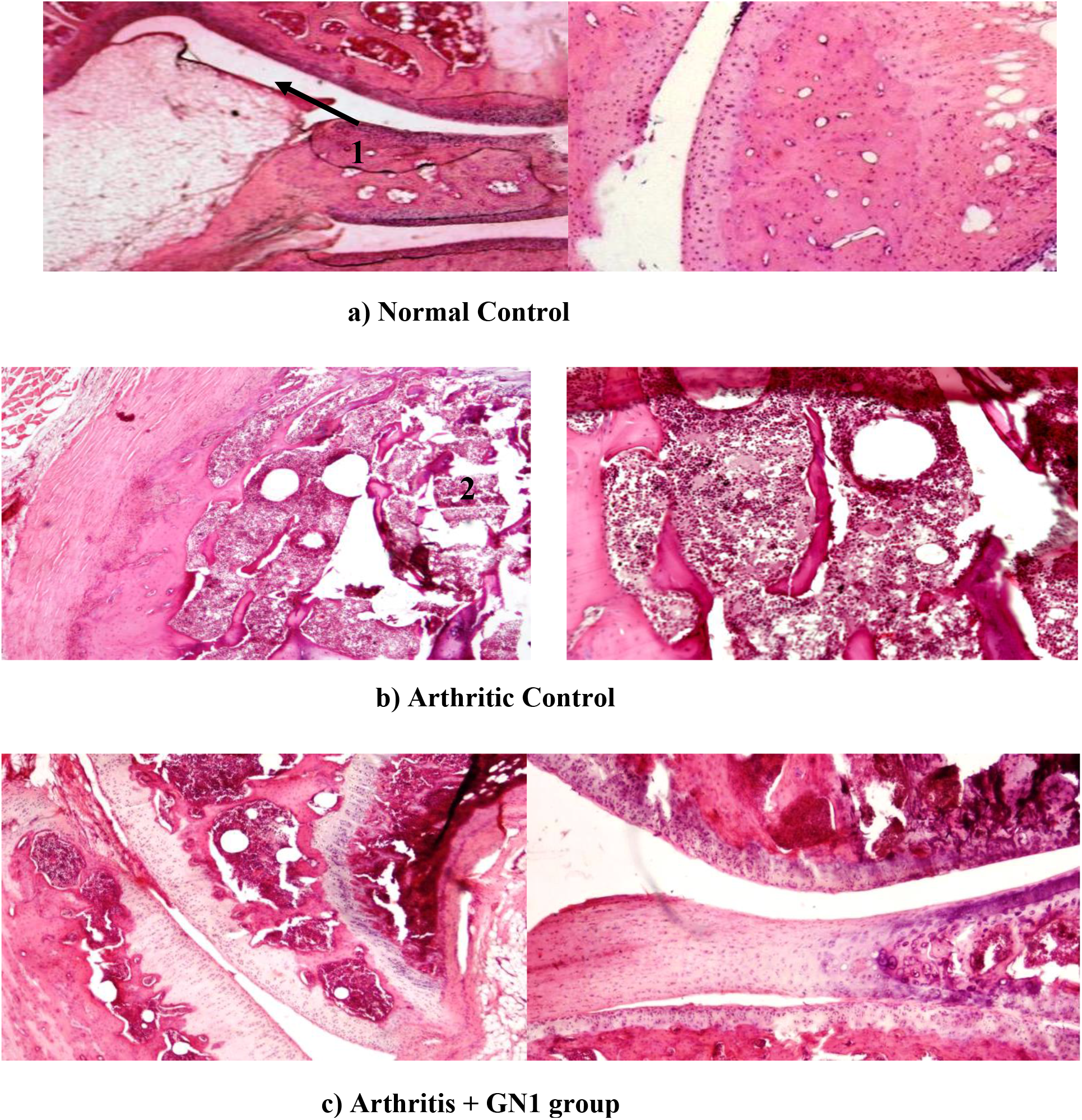
Photomicrographs showing H & E stained knee joints from a normal, arthritic control rats and GN1 treated arthritic rats. A prominent lymphocytic infiltration of synovium with invasion of periarticular bone, collapse of articular surface and articular bone destruction and pannus formation over articular cartilage can be seen in arthritic group. In GN1 treated animal joint histological studies shows intact articular cartilage with mild infiltration of articular surface with slight periarticular tissue destruction.

The concentration of IL-1β and TNF-α was determined using ELISA assay kit in the serum (day 22) of normal control, arthritic rats, arthritic rats treated with indomethacin, GN1. The results (Figure 6 and 7) show that in arthritic rats the level of TNF α (P < 0.001) and IL-1 β (P < 0.005) were significantly increased as compared to non-arthritic control rats. While indomethacin was able to significantly inhibit the concentration of circulatory TNF-α (P < 0.001) as compared to arthritic rats and it also significantly decreases the serum IL-1 ß level (P < 0.002). It was also found that both pro-inflammatory cytokines level i.e., TNFα and IL-1β were significantly in the arthritic rats treated with GN1 (P < 0.005, P < 0.015 respectively) as compared to untreated arthritic rats.

**FIGURE 6:**
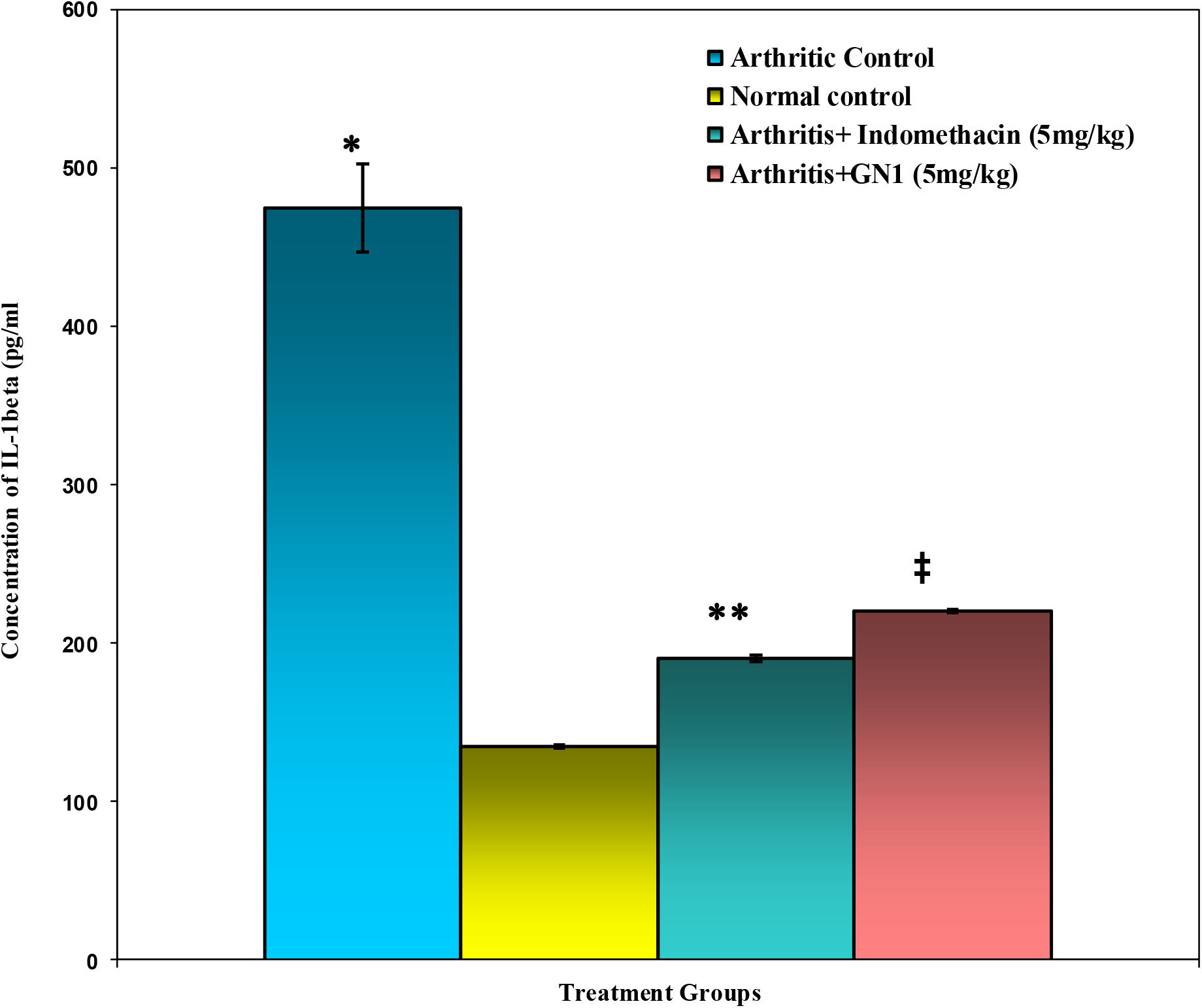
The effect of GN1 (5 mg/kg) on the concentration of IL-1ß (pg/ml) measured in the serum of arthritic and non-arthritic rats. Data is represented as means ± SEM for 12 animals in each group. The serum level of IL-1ß was found significantly higher in the arthritic control group compared to normal animals (*p < 0.001). In contrast, both the treatment of indomethacin and GN1 were found to decrease the level of the measured IL-1ß compared to the untreated arthritic group (where **p < 0.009, ^‡^p < 0.005, for indomethacin-treated and GN1 treated group respectively).

**FIGURE 7:**
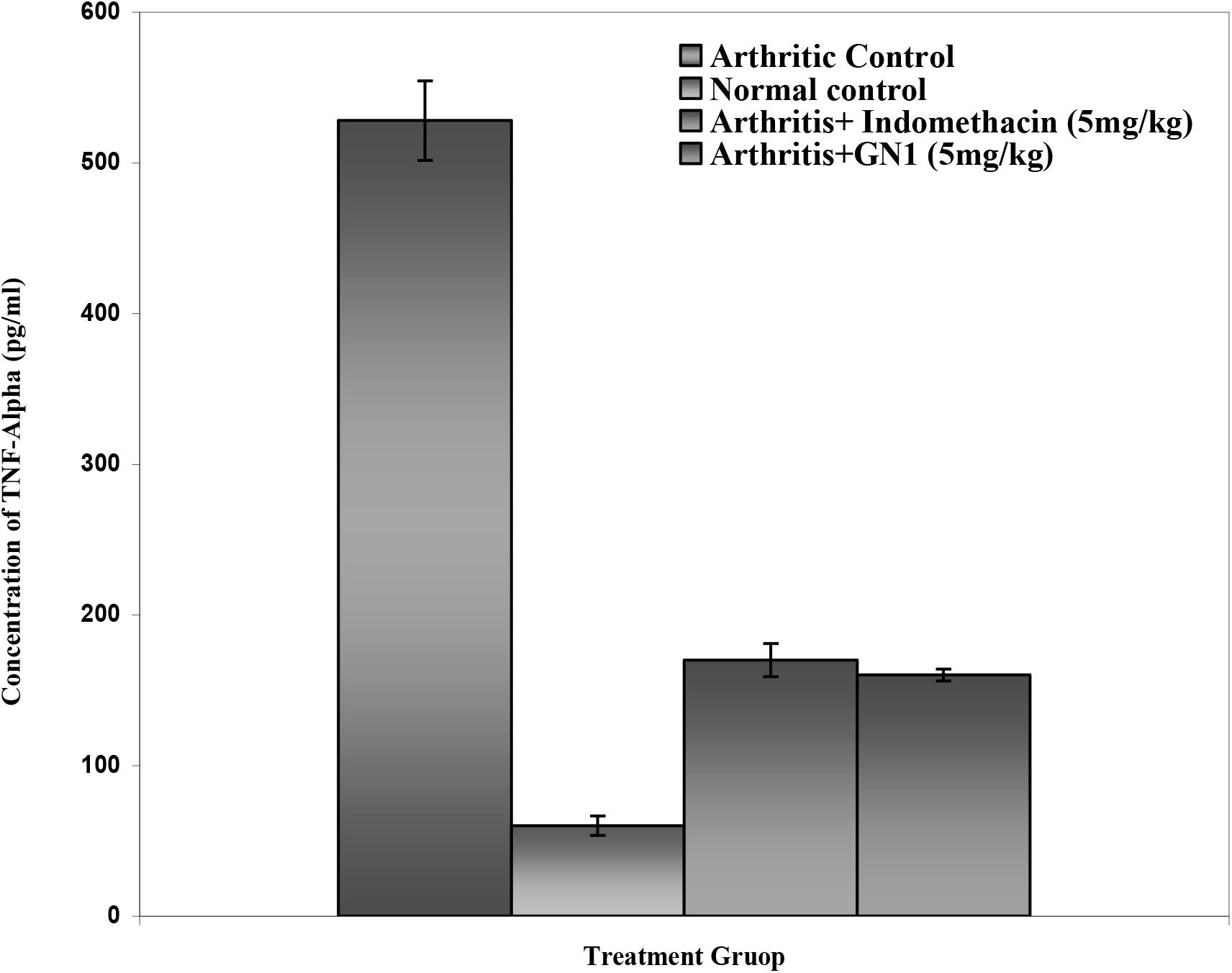
The concentration of TNF-α (pg/ml) measured in the serum of arthritic and normal rats. Each bar represents the means ± SEM for n = 12 animals/group. The arthritic control group exhibited a significant increase in the level of TNF-α (*p < 0.0009) in comparison to normal control group. In contrast, the treatment of GN1 also demonstrated a marked reduction in the measured parameter (**p < 0.015). Likewise, the arthritic group receiving the treatment of indomethacin (5 mg/kg) showed a significant reduction in the serum TNF α (^‡^p < 0.001).

The concentration of glutathione (GSH), nitric oxide (NO) and peroxide were determined by colorimetric method in the serum (day 22) of the experimental groups (Table 1). There was a marked significant decreased in the concentration of GSH (P < 0.021), and significant increase in the formation of NO (P <0.03) and peroxide (P < 0.001) in the arthritic control rats as compared to non-arthritic control rats. GN1 treatment in arthritic rats was significantly increase the GSH level (P <0.037) and was capable to decrease the production of peroxide significantly (P < 0.002). However, GN1 can suppress the production of nitric oxide in arthritic rats, but this inhibition was not as much as significant. The treatment of reference drug indomethacin did not show any effect on the GSH and nitric oxide level in the arthritic rats as compared to untreated arthritic rats. While, it is able to decrease the production of peroxide significantly (P < 0.038) as compare to arthritic control rats.

**TABLE 1:** Effect of GN-1 (5 mg/kg) on the oxidative stress markers following the induction of arthritis. Each value represents the mean ± SEM of 12 animals per group.

| Treatment Groups | OXIDATIVE STRESS MARKERS |  |  |
| --- | --- | --- | --- |
|  | GSH (mg/dl) | NO (mg/dl) | PO (ng/dl) |
| Normal Control | 20.624 ± 0.7 | 0.023 ± 0.001 | 273.7 ± 14.04 |
| Arthritic Control | 8.416 ± 0.610 | 0.102 ± 0.004 | 745.5 ± 14 |
| Indomethacin (5 mg/kg) + Arthritis | 13.848 ± 0.5 | 0.0465 ± 0.006 | 572 ± 14.049 |
| GN-1 (5 mg/kg) + Arthritis | 18.392 ± 0.6 | 0.042 ± 0.0042 | 420 ± 14.05 |

**Table 2:**
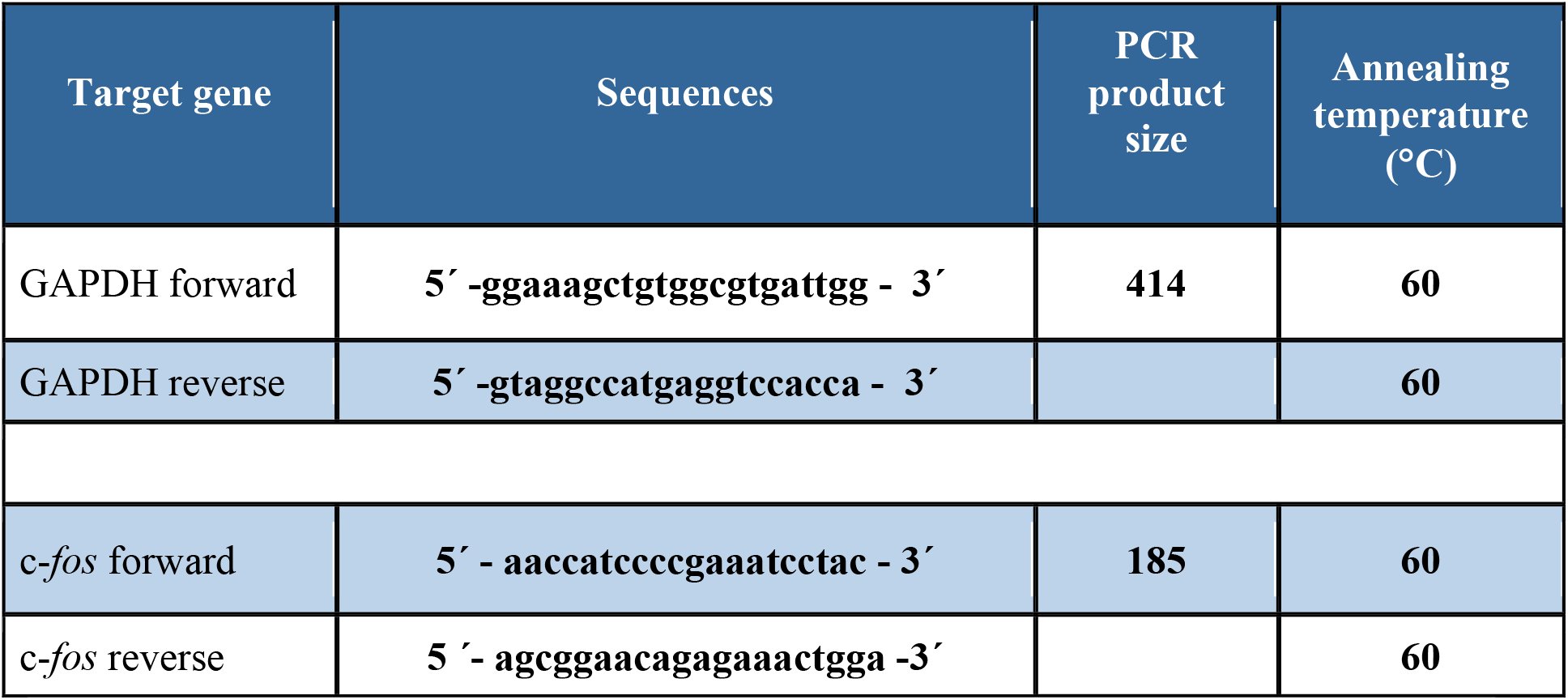
Primer Sequences of GAPDH and *cfos* gene.

Since exposure to variety of acute stress stimuli including acute immune challenge has been reported to increase the *c-fos* mRNA, therefore we have used the *c-fos* mRNA within the central nervous system as a marker of neuronal activation following arthritis-induce immune stress. We observed baseline expression or constitutive expression of c-*fos* in normal control animals, however, when statistically analyzed it was found to be non-significant (fig. 8 and 9).

**Figure 8:**
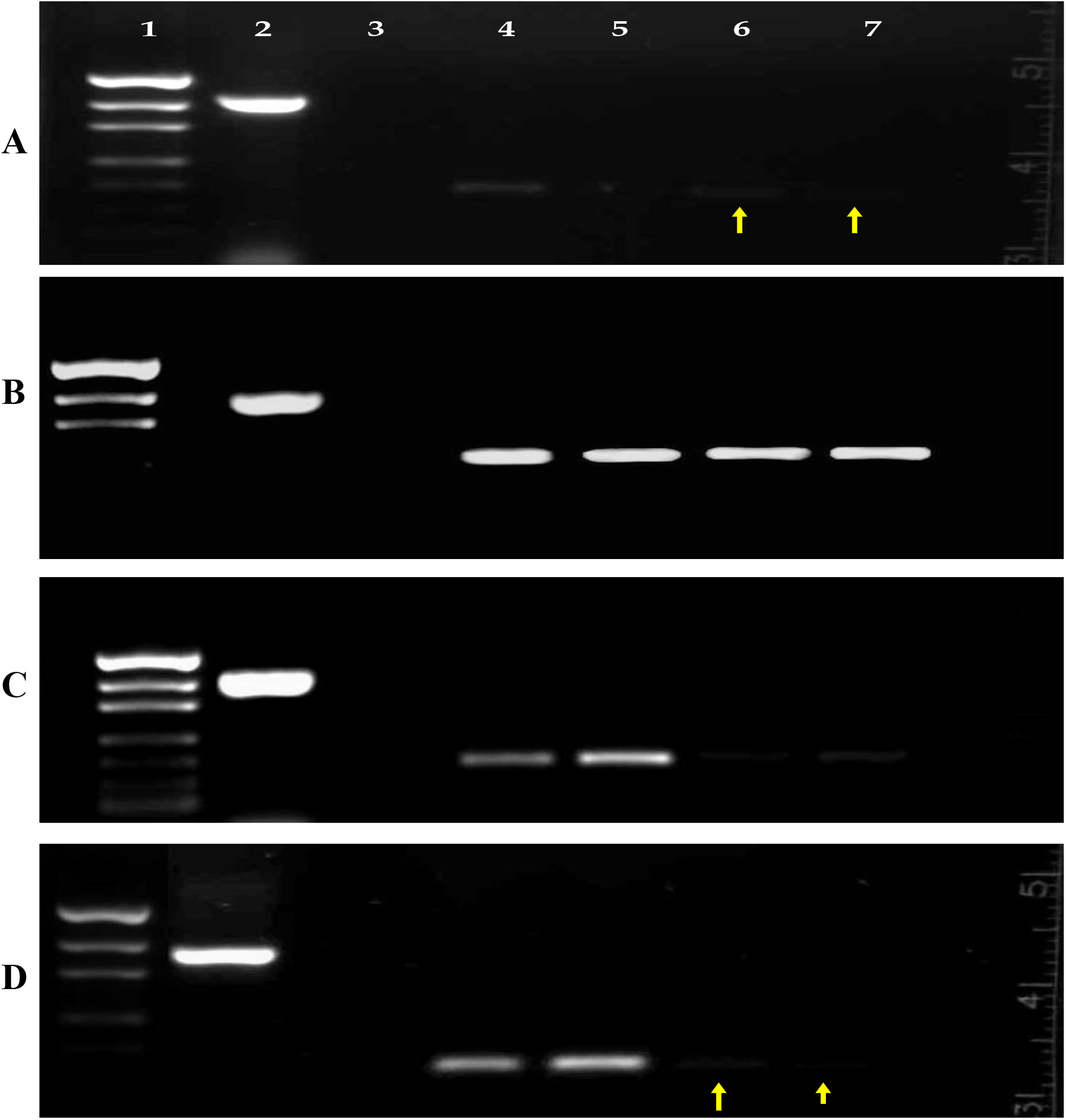
Effects of GN1 on c-*fos* mRNA level. Total RNA was isolated and subjected to RT-PCR analysis and finally the PCR products were visualized on 1% agarose gel. Gel A, B, C and D represents normal, arthritic control, Indomethacin and GN1 treated groups respectively. The lanes 1-7 represents DNA molecular weight marker, GAPDH (414 bp), RT negative, and expression of c*-fos* in hippocampus, thalamus, cortex and amygdala, respectively. The PCR product was quantified by densitometry and each c-*fos* mRNA level was normalized to that for GAPDH mRNA.

**Figure 9:**
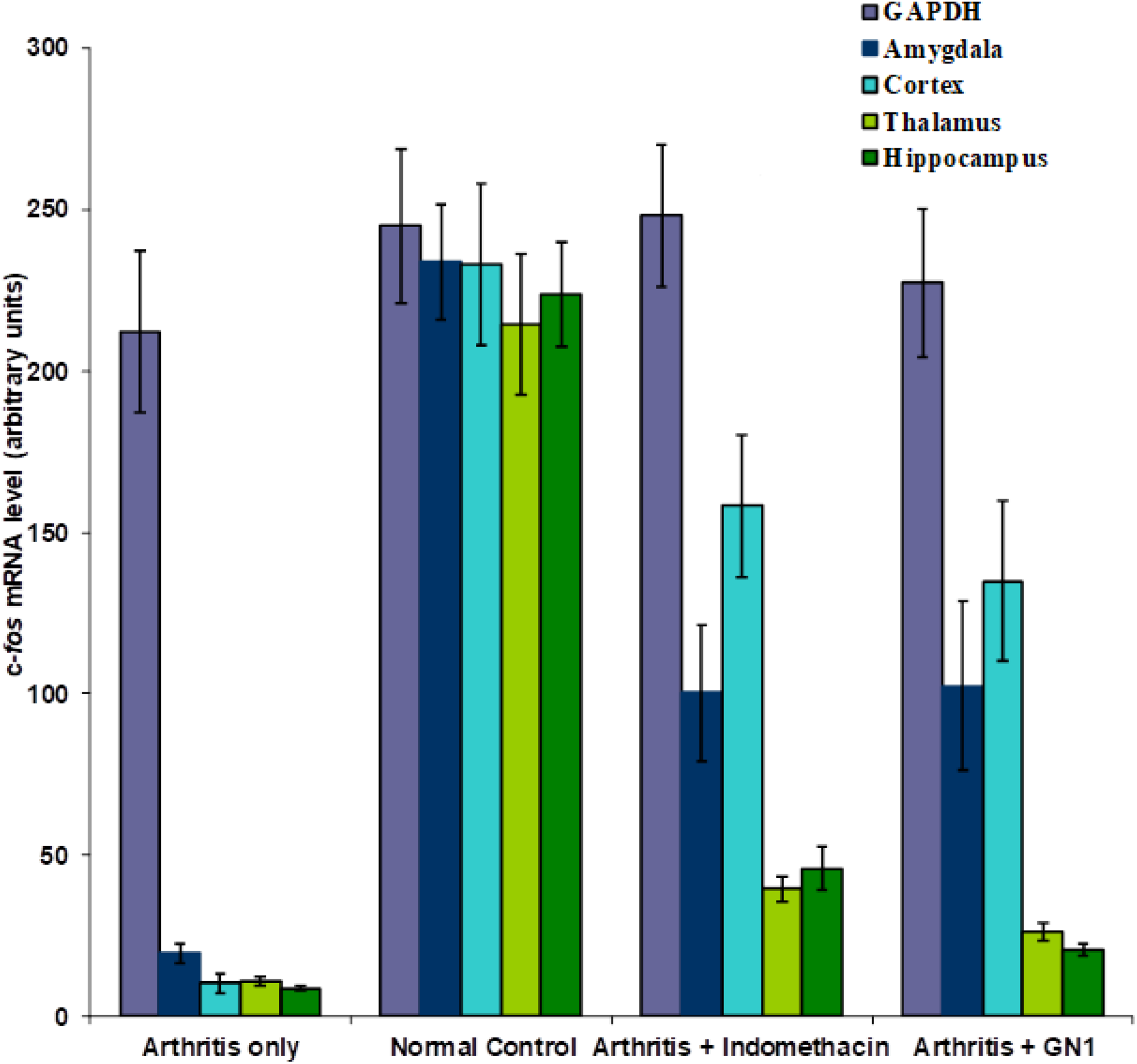
Effect of GN1 on *c-fos* mRNA levels. Following the agarose gel electrophoresis, the bands were quantified by densitometry and each c-*fos* mRNA level was normalized to that for GAPDH mRNA. Each bar represents the mean ± S.E.M.

The RT-PCR reactions were preceded for the specific band of *c-fos* gene (185 bp) in four different regions i.e., amaygdala, cortex, hippocampus and thalamus of the brain samples from treated, untreated arthritic and normal control groups (fig. 8). A densitometry analysis revealed a baseline expression of *c-fos* in amygdale, thalamus and hippocampus regions of brain samples from normal control animals (fig. 9). In all experiment, the expression of housekeeping GAPDH gene (414 bp) was also analyzed and it was found that to be equal among groups i.e., normal control, treated and untreated arthritic control brain samples. When arthritic control samples were compared with the normal control group, a marked increase in the mRNA expression of *c-fos*. Through densitometry analysis, amongst the various regions tested, the highest concentration of *c-fos* mRNA was detected in the amygdala and cortex followed by hippocampus and finally in the thalamus. Next, we compared the drug control i.e., indomethacin treated arthritic rats with that of arthritic control group and observed a reduced *c-fos* mRNA expression which was maximum in the thalamus followed by hippocampus, then amygdala and finally in cortex. Densitometry analysis performed on the PCR products from various treatment groups demonstrated that all the treatment modalities markedly reduce the expression of *c-fos* mRNA and maximum reduction in all cases was observed in the thalamus followed by hippocampus. Although the cortical regions and amygdala of the brain samples from GN-1 treated groups also demonstrated marked reduction in *c-fos* expression, however, compared to thalamus and hippocampus it was found to be slightly more.

Results and observations lead to the hypothesis that the joint inflammation modifying abilities of the GN1 would reduce the development and severity of the arthritis. Biochemical studies revealed that GN1 can inhibit the production of pro-inflammatory marker c-fos, cytokines such as TNF-α and IL-1β which is the most primarily mediator of certain inflammatory responses. It also maintained the body redox state by keeping the level of GSH as same as in non-arthritic rats. Furthermore, it also helps in regulate the production of nitric oxide and peroxide formation which in turn inhibits the stimulation of various inflammatory cascades such as infiltration of lymphocytes and leukocytes into the joint. These above effects of the GN1 in the inflammatory environment may due its anti-inflammatory and immunosuppressive actions.

Finally we can hypothesized that GN1 suppresses the progression of disease in the AIA rat models, a model of RA, by inhibiting the inflammatory reaction such as suppression the of nitric oxide and peroxide formation, down regulation of TNF-α, IL-1β and c*-fos* etc, suggesting a possible therapeutic potential in the treatment of the disease.

